# Disruption of a structural niche for otoconia maintenance may underlie common vestibular disorders

**DOI:** 10.64898/2026.09.16.752180

**Authors:** Diana M. Correa, Abel P. David, Ruiqi Zhou, Richard Osgood, Artur A. Indzhykulian, Stephan Krämer, Taha A. Jan, Andreas H. Eckhard

**Affiliations:** Otopathology Laboratory, Mass Eye and Ear, Boston, MA, USA; Eaton-Peabody Laboratories, Mass Eye and Ear, Boston, MA, USA; Department of Otolaryngology-Head and Neck Surgery, Harvard Medical School, Boston, MA, USA; Department of Otolaryngology-Head and Neck Surgery, Northwestern Medicine, Chicago, IL, USA; Feinberg School of Medicine, Northwestern University, Chicago, IL, USA; Department of Otolaryngology-Head and Neck Surgery, Epithelial Biology Center, Vanderbilt Center for Stem Cell Biology, Center for Computational Systems Biology, Vanderbilt University Medical Center, Nashville, TN, USA; Center for Nanoscale Systems, Harvard University, Cambridge, MA, USA

**Keywords:** otolith organs, vestibular system, benign paroxysmal positional vertigo, endolymphatic hydrops, extracellular matrix, epithelial-mesenchymal signaling, falls

## Abstract

Many falls and balance disorders in older adults originate in the otolith organs, the gravity sensors of the inner ear. These sensors maintain upright posture through otoconia, calcium carbonate crystals that mass-load the sensory maculae. Otoconia dislodgement causes the most common form of vertigo, and their age-related loss reduces gravity sensation and undermines balance. Yet the cellular mechanisms of otoconia formation and maintenance—and how they fail in disease— remain unknown. Using mineral-preserving histology, crystal-sensitive imaging, volume electron microscopy, and immunolabeling in human and animal otolith organs, we found otoconia biogenesis–related proteins and early crystallization at the pole opposing the macula, the roof domain. We discovered filigree extracellular matrix scaffolds bridging roof and macula, loaded with nascent otoconia, suggesting scaffold-guided transport across the organ. Single-cell transcriptomics nominated a specialized roof mesenchyme signaling to the roof epithelium as a driver of otoconia and scaffold production. In guinea pigs with endolymphatic hydrops, fluid expansion of the organs ruptured the otoconia-trafficking scaffolds as roof and macula separated, followed by a decline in macular otoconial mass. We propose a new disease model for common vertigo and balance disorders in which disruption of the otoconia-generating and -trafficking epithelial-mesenchymal roof niche leads to displacement and depletion of otoconia.

## Introduction

Falls and chronic imbalance are a major public health burden in aging populations (1, 2), where more than one quarter of adults aged 65 or older fall each year (3). Fall-related mortality rises sharply with age (3, 4), and approximately 684,000 people die from falls worldwide each year (3). Among vestibular causes of falls, imbalance, and dizziness, the condition of benign paroxysmal positional vertigo (BPPV) is the most common diagnosis (5–7), classically triggered by lying down, turning in bed, or other head movements (8). The inner ear’s two sac-like gravity-sensing otolith organs are the utricle and the saccule, which contain otoconia that facilitate gravity detection. Otoconia are micrometer-scale calcium carbonate crystals that overlie the sensory maculae, and their mass loads the maculae so that gravity’s pull and linear acceleration register in the horizontal (utricle) and vertical (saccule) planes (9–12). In BPPV, otoconia are dislodged from the utricular macula and enter a semicircular canal, where head turns set off brief, intense spinning vertigo, often accompanied by severe nausea and unsteadiness (5, 13–15). With age, macular otoconia degenerate and decline in number (16, 17), gravity sensing weakens (12), and this loss of vestibular function contributes to the imbalance and falls of later life (1).

Yet basic questions remain unanswered: how are otoconia formed and maintained, and how do these processes fail in vestibular disease? In mice, zebrafish, and birds, the proteins building otoconia and otoliths are produced mainly by the non-sensory roof domain, the epithelium facing the macula across the endolymph-filled lumen, rather than the otoconia-loaded macula itself (18–20, reviewed in ref. 21). Otoconia thus appear to originate not at the site where they are ultimately found and function, but on the opposite wall of the organ. It has never been established where these proteins mineralize, how otoconia then cross the lumen from the roof to the macula, or how they are precisely patterned on the macular surface—nor is it known whether the same arrangement holds in the human otolith organs. These fundamental gaps have clinical consequences. Current models of BPPV (5, 6) and of age-related otoconia decline (16, 17) begin with mature otoconia already on the macula and describe downstream processes, including cracking, demineralization, and detachment, but neglect the processes that produce them. These models, therefore, can neither explain why otoconia detach or decline nor indicate how loss might be prevented.

Here we address these questions in human and animal otolith organs, combining tissue-preserving methods we developed (22) to keep otoconia and the luminal architecture intact *in situ*, relationships that are destroyed with standard histological processing (23). We use volume electron microscopy to resolve otoconia at their earliest stages of mineralization, and single-cell RNA sequencing (scRNA-seq) to identify the cellular origins of otoconia biogenesis. We show that the roof domain is a conserved epithelial-mesenchymal niche in which otoconia begin to form, and that previously unrecognized extracellular matrix (ECM) scaffolds link this niche to the maculae and carry nascent otoconia. In a pathologic experimental model of endolymphatic hydrops, which distends the lumen and separates roof from macula, the scaffolds ruptured and led to a decline in macular otoconial mass. Together, these findings identify a niche that supplies otoconia throughout adult life and provide structure-based models for their interrupted trafficking and loss.

## Results

### In mouse otolith organs, nascent otoconia populate the roof domain and filigree ECM scaffolds that bridge the lumen from roof to macula

The non-sensory roof domain of the otolith organs has previously been implicated in otoconia formation (20, 21). We therefore focused on the adult mouse otolith organs prepared in two complementary ways: decalcified, celloidin-embedded sections, which retain only the organic (protein) matrix of otoconia, and undecalcified (mineralized), methyl methacrylate resin– embedded sections, which retain mineral and matrix *in situ*. In both preparations, the roof domain contained nascent, crystallizing otoconia, consistent with roof-localized production of otoconial proteins (20, 21). Unexpectedly, the roof domain was also connected to the macula by a previously unrecognized ECM scaffold spanning the lumen. In the saccule, this scaffold was a slender conduit running from the anterior roof domain to the anterior macular margin (Figure 1A). It was glycoprotein rich (periodic acid–Schiff [PAS] positive), filamentous, and continuous with the macular otoconial layer, and carried discrete otoconial packets along its length (Figure 1B). The nascent otoconia-containing scaffold was readily distinguished from artifactually dislodged otoconial membranes (Supplemental Figure 1). Composite mapping of 26 saccules with serial-section three-dimensional (3D) reconstruction demonstrated that the conduit was consistently present and stereotyped in course, with a dominant anterior macular insertion and a smaller, variable posterior branch (Figure 1, C and D). This conduit was fibronectin positive and contained otoconin-90 (Oc90)-positive particles (Figure 1, E and F). In mineralized preparations, polarized light revealed birefringent crystals within the scaffold and around its anterior macular insertion (Figure 1, G and H), suggesting that roof-derived otoconial material begins mineralizing before reaching the macula. *En face* macula morphometry reconstructed from serial sections showed the smallest crystals at the conduit’s anterior insertion and progressively larger, mature otoconia toward the posterior macular margin (Figure 1, I and J). The maturation gradient begins where the scaffold meets the macula, consistent with maturing otoconia advancing across the macula.

**Figure 1.**
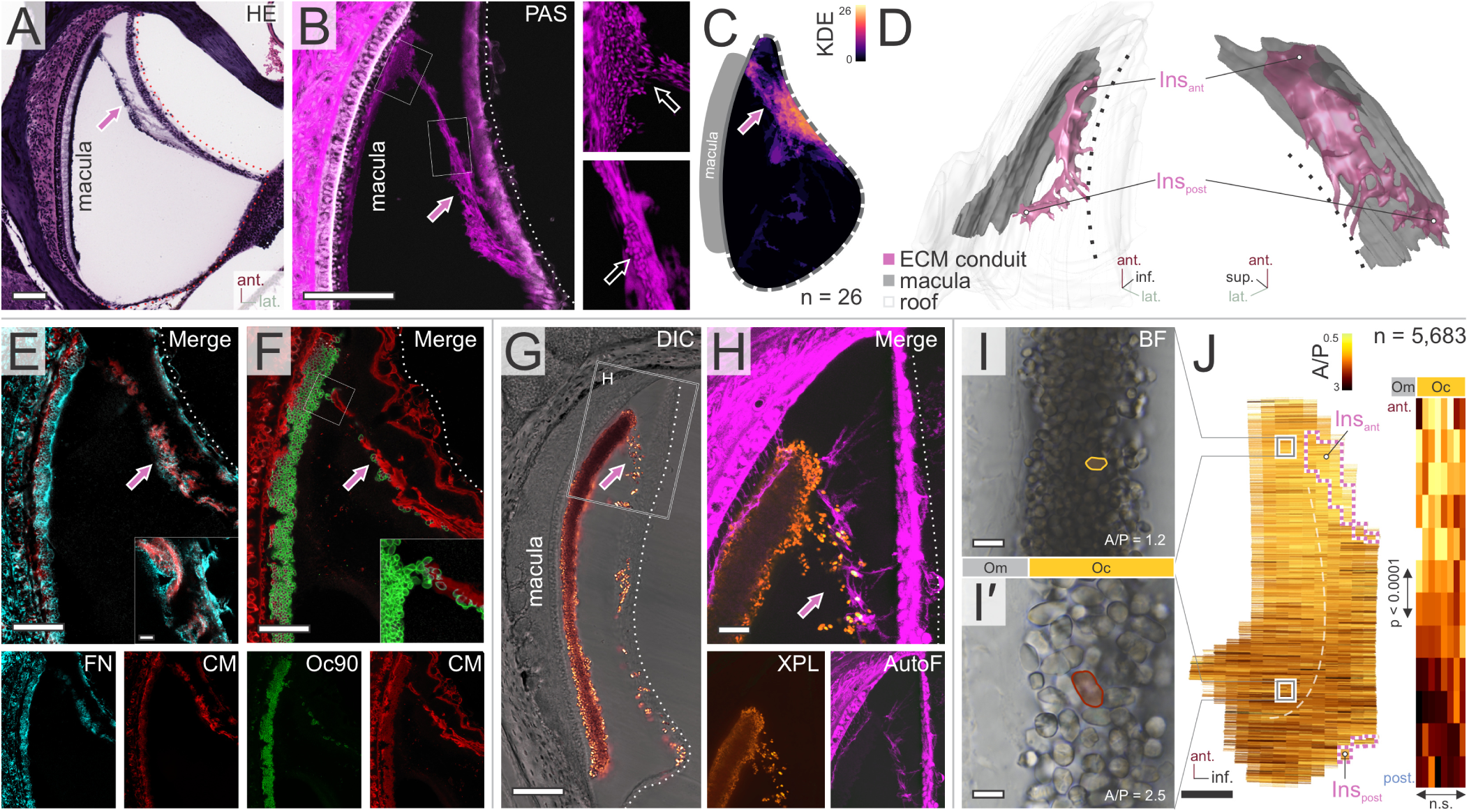
A roof-derived extracellular matrix conduit connects the roof domain to the macula and carries nascent otoconia in the mouse saccule. (**A**) Decalcified, celloidin-embedded saccule, hematoxylin and eosin (H&E): otoconial conduit (arrow) from roof domain (dotted line) to anterior macular margin. (**B**) Same preparation, confocal periodic acid–Schiff (PAS) fluorescence: glycoprotein-rich conduit (arrow) from the roof (dotted line); insets, anterior macular insertion and otoconia-like particles within the conduit. (**C**) Segmented conduits of 26 saccules overlaid as a kernel density estimate (KDE) map: stereotyped roof-macula trajectory (arrow). (**D**) 3D reconstruction of a serially sectioned saccule: anterior (Ins_ant) and posterior (Ins_post) conduit insertions; dotted line, roof. (**E**) Decalcified celloidin section, fibronectin immunofluorescence (cyan) with CellMask Deep Red (CM; red): fibronectin-positive, CM-labeled scaffold (arrow; inset, higher magnification). (**F**) Otoconin-90 (Oc90; green) and CM: Oc90-positive particles within the conduit, merging into the macular otoconial layer at its anterior insertion. (**G**) Mineralized, Technovit-embedded saccule, differential interference contrast (DIC): sail-like scaffold (arrow); box, region in **H**. (**H**) Cross-polarized light (XPL) with autofluorescence (AutoF): birefringent otoconia within the scaffold, continuous with the macular layer. (**I** and **I**′) Brightfield, mineralized section: macular otoconia near the anterior (**I**) and posterior (**I**′) insertion; one otoconium outlined (yellow) with area-to-perimeter (A/P) ratio. (**J**) Reconstruction from PAS-stained serial sections (confocal), area and perimeter of every otoconium: en face heat map (white to dark red-brown) and depth view of mean A/P ratio; anterior-to-posterior gradient (P < 0.0001, linear regression); Oc/Om, otoconia/otoconial membrane (n.s., not significant). Magenta boxes, anterior and posterior insertion points of the conduit (Ins_ant and Ins_post in D). n = 5,683 crystals from 1 serially sectioned saccule. Scale bars: 100 μm (A, B, and G); 50 μm (E and F); 20 μm (H); 10 μm (I and I′); 200 μm (J).

A similar pattern is seen in the utricle, where the roof domain opposite the macula comprises a specialized epithelium called the vestibular dark cell (VDC) epithelium. From the VDC, as from the saccular roof, arose an ECM scaffold, in this case consisting of a broader, diffuse otoconial web overlying the entire macula, and a narrower medial branch directed toward an adjacent VDC field near the horizontal semicircular canal crista (Figure 2, A–C). These structures were likewise glycoprotein and proteoglycan rich (PAS-Alcian blue positive), reproducibly positioned relative to the macula by composite mapping and 3D reconstruction (Figure 2, D and E), and enriched for fibronectin and Oc90 (Figure 2, F and G). Mineralized preparations again localized otoconial material within and along the scaffold, including the medial branch at the macular margin (Figure 2, H and I). Macular otoconia were again graded in size, though along a different axis: utricular otoconia varied little from medial to lateral but showed a prominent depth gradient within the otoconial layer, smallest toward the lumen (Figure 2, J and K), consistent with seeding across the whole macular surface followed by maturation in place.

**Figure 2.**
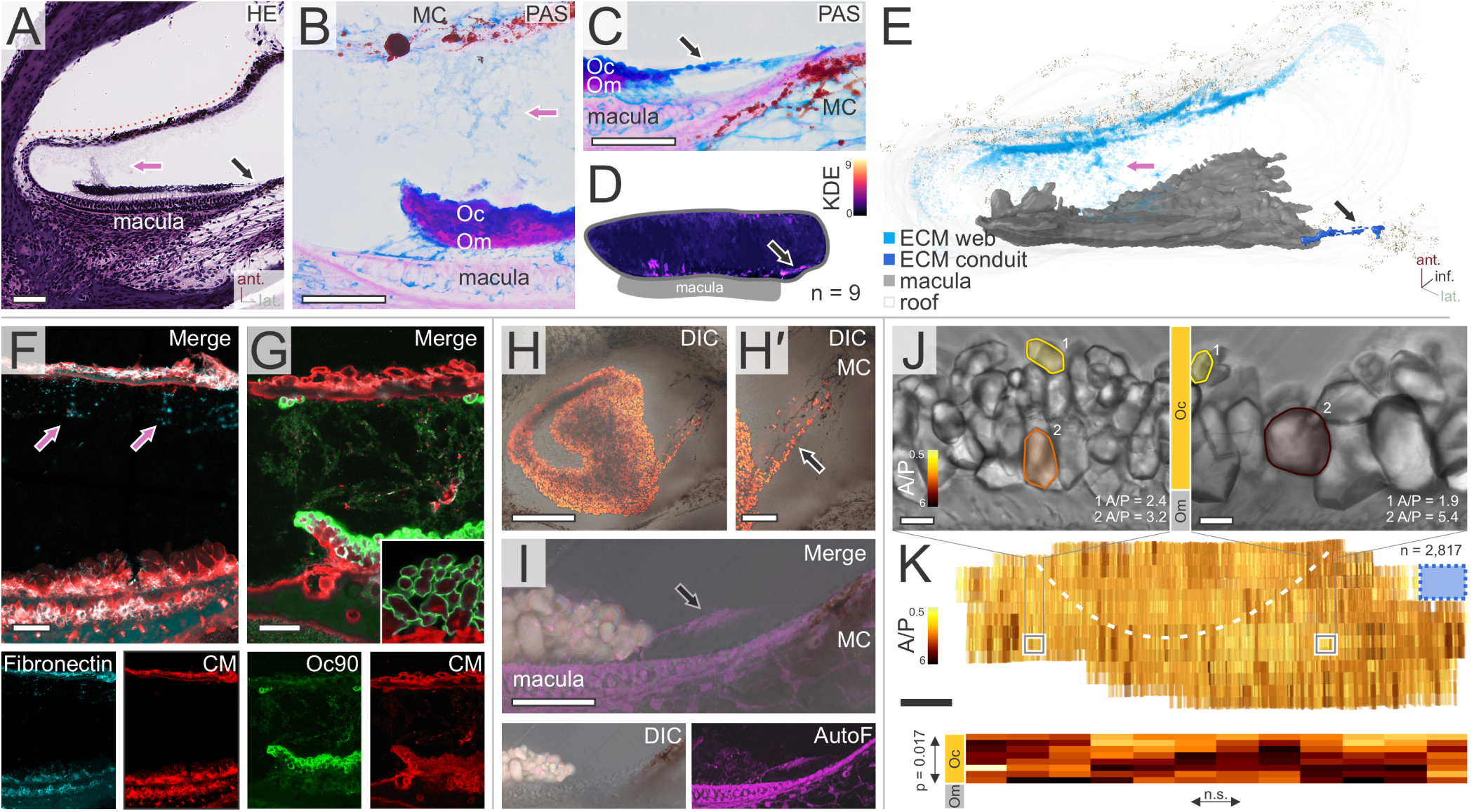
A roof-derived extracellular matrix (ECM) web connects the roof domain to the macula and carries otoconial material in the mouse utricle. (**A**) Decalcified, celloidin-embedded utricle, hematoxylin and eosin (H&E): web (magenta arrow) from roof (dotted line) to macula; medial branch (black arrow) from the horizontal canal crista region to the lateral macular margin. (**B**) Periodic acid–Schiff (PAS)-Alcian blue, high power: Alcian blue–positive web (magenta arrow); Oc, otoconial layer; Om, otoconial membrane; MC, melanin-containing cells. (**C**) PAS-Alcian blue: medial branch (arrowhead) with Alcian blue–positive otoconia-like particles. (**D**) Tracings of 9 utricles as a kernel density estimate (KDE) map: diffuse web, consistent medial branch (arrow). (**E**) 3D reconstruction: web (cyan; magenta arrow), medial branch (blue; black arrow), macula (dark gray), roof (light gray). (**F**) Decalcified celloidin section, fibronectin immunofluorescence (cyan) with CellMask Deep Red (CM; red): punctate fibronectin along the web near the roof. (**G**) Otoconin-90 (Oc90; green) and CM: Oc90-positive particles at the roof, diffuse labeling of the web across the lumen; inset, circumferentially labeled macular otoconia. (**H** and **H**′) Mineralized, Technovit-embedded utricle, en face differential interference contrast (DIC): otoconial layer (**H**); medial branch loaded with small otoconia (arrowhead, **H**′). (**I**) DIC with autofluorescence (AutoF): medial branch from the MC zone to the lateral macular margin. (**J**) Brightfield: macular otoconia with area-to-perimeter (A/P) ratios, smaller near the endolymph (Oc), larger near the otoconial membrane (Om). (**K**) Reconstruction from PAS-stained serial sections (confocal), A/P ratio of every otoconium: no gradient across the macula en face (n.s.); depth view, gradient from endolymphatic to macular side (P = 0.017, linear regression); boxes, fields in **J**; dashed line, striola; blue box, site where the medial branch from the VDC field near the horizontal canal crista inserts into the otoconial membrane. n = 2,817 crystals from 1 serially sectioned utricle. Scale bars: 100 μm (A); 50 μm (B, C, F, G, H′, and I); 200 μm (H and K); 10 μm (J).

In both otolith organs, therefore, the roof domain gives rise to an ECM scaffold that physically couples it to the macula and carries otoconia at successive stages of mineralization. Between the two organs, the scaffolds differ in form: a discrete conduit in the saccule versus a diffuse web in the utricle. In both organs, crystal size appears to form a gradient that originates from the point of contact of the trafficking scaffold with the macula. This arrangement raised three questions, which we addressed next: 1) whether these ECM scaffolds are static developmental remnants or dynamic structures that continuously maintain the otoconial mass in adult life; 2) whether the same arrangement exists in the otolith organs of humans or other vertebrates; and 3) which transcriptional programs and signaling pathways equip the roof domains to produce otoconial matrix and scaffold.

### Disruption of roof-macula ECM scaffolds in adult otolith organs is followed by progressive loss of the macular otoconial mass

To test whether the scaffolds maintain the adult otoconial mass, we disrupted them in adult animals and monitored the otoconial layer afterward. Since the scaffolds span the lumen and cannot be reached surgically without destroying the organ, we disrupted them indirectly, by enlarging the lumen instead. We used a well-characterized guinea pig model of endolymphatic hydrops, where the endolymphatic duct, which connects the inner ear’s fluid space to the sac that maintains endolymph volume and composition, is mechanically obstructed. This obstruction results in the progressive expansion of the lumen (endolymphatic hydrops) (24, 25). As the lumen expands, the roof and macular epithelia grow further apart and the scaffolds spanning between them must lengthen, a strain we predicted would rupture them (Figure 3, A and F). In archival material from operated animals, the endolymphatic volume of both organs, the histological measure of hydrops, was increased at every survival time, rising most steeply between 2 and 8 weeks and more in the saccule than in the utricle (Figure 3, B and G), consistent with prior reports (24, 26). In control animals, the saccular conduit was intact and both maculae carried a dense, continuous otoconial layer (Figure 3, C, E, H, and J). In animals with endolymphatic hydrops at 2 weeks, distension was visible but modest, the conduit remained continuous, and neither otoconial layer differed from controls (Figure 3, E and J). By 8 weeks both organs were markedly distended. In the saccule, the conduit had ruptured, leaving fragments at the roof domain and the anterior macular insertion, and the otoconial layer had lost its compact organization and dispersed into the lumen (Figure 3, D and E). In the utricle, where the web could not be reliably visualized but the roof-macula distance had increased markedly, the otoconial layer had only begun to thin (Figure 3, I and J). By 16 weeks, the saccular macula had lost its otoconial layer entirely, and the utricular layer was reduced to a sparse remnant (Figure 3, E and J). These findings establish a close temporal relationship between scaffold rupture and progressive loss of the macular otoconial layers, consistent with scaffolds that maintain the adult otoconial mass rather than persisting as developmental remnants.

**Figure 3.**
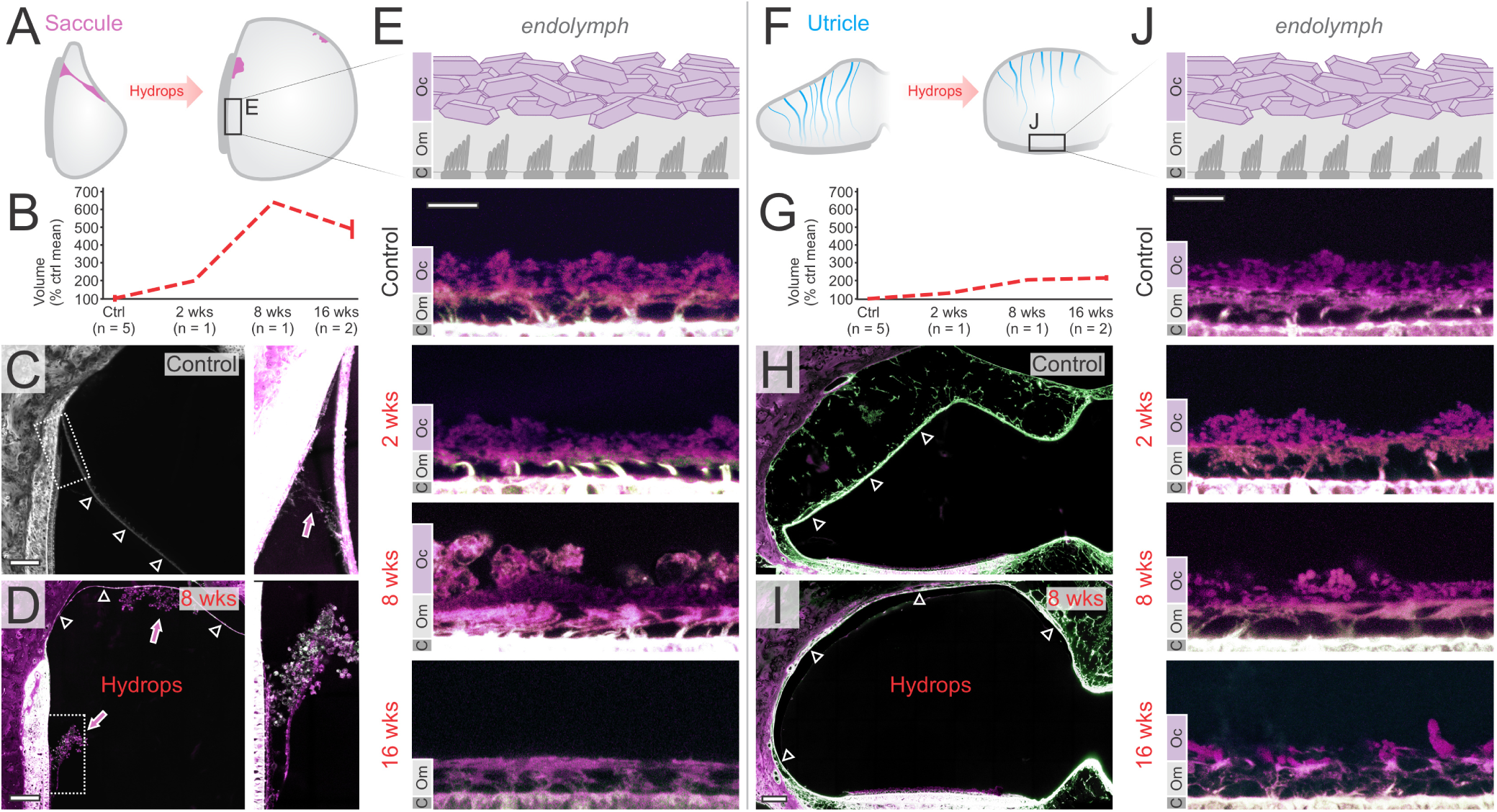
Experimental endolymphatic hydrops disrupts the roof-macula extracellular matrix (ECM) scaffold before depleting the macular otoconial layer in the guinea pig. All micrographs: confocal eosin fluorescence (green) and tissue autofluorescence (magenta), decalcified celloidin sections (otoconia represented by their organic matrix). (**A**) Saccule schematic: ECM conduit (magenta) between roof and macula; with hydrops, the roof distends away from the macula and the conduit ruptures; box, region in **E**. (**B**) Saccular endolymphatic volume (percentage of control mean; mean ± SD where n > 1): controls and 2, 8, and 16 weeks after surgery (n = 5, 1, 1, and 2 ears). (**C** and **D**) Control saccule (**C**): roof (arrowheads) and conduit (magenta arrow) to the anterior macular margin; 8 weeks (**D**): roof (arrowheads) displaced, conduit remnants on macula and roof (magenta arrows); boxed regions enlarged at right. (**E**) Saccular macula: schematic above (Oc, otoconial layer; Om, otoconial membrane; **C**, apical surface of the sensory epithelium) and confocal series below, control and 2, 8, and 16 weeks: otoconial layer unchanged at 2 weeks, markedly thinned by 8 weeks, lost by 16 weeks. (**F**) Utricle schematic: ECM web (blue) thins and detaches from the macula with hydrops; box, region in **J**. (**G**) Utricular endolymphatic volume, same ears. (**H** and **I**) Control utricle (**H**): roof (arrowheads) near the macula; 8 weeks (**I**): roof (arrowheads) displaced against the bony wall. (**J**) Utricular macula, same layout and time course as E: preserved at 2 weeks, then progressively depleted to a sparse remnant by 16 weeks. Group sizes preclude statistical comparison; **B** and **G** are descriptive. Scale bars: 100 μm (**C**, **D**, **H**, and **I**); 10 μm (**E and J**).

### In human otolith organs, otoconia biogenesis begins inside roof epithelial cells and proceeds along ECM scaffolds toward the macula

The same roof-macula arrangement was present in adult human otolith organs, examined in decalcified and mineralized preparations. In the human saccule, the anterior roof domain is known to be morphologically specialized, with a thickened subepithelial, perilymph-facing mesenchyme—the reinforced anterior roof area (RARA) (23). From this domain, in the same position as in mouse and guinea pig, a slender ECM conduit again extended toward the anterior macular surface (Figure 4, A and B), and otoconia-like protein remnants in decalcified material (Figure 4A′) and birefringent crystals in mineralized material (Figure 4, B and B′) again marked nascent otoconial packets within it. Within the RARA, particulate structures occurred both inside epithelial cells and in the subepithelial space, ranging from weakly birefringent ovoid particles (Figure 4C) to fully birefringent otoconia (Figure 4C′); comparable structures were absent from roof epithelium outside the RARA. Focused ion beam–scanning electron microscopy (FIB-SEM) of the RARA in an adult human saccule resolved matrix-dense domains within roof epithelial cells, consistent with intracellular crystallization (Figure 4, E and E′), and ECM strands extending from the apical membrane into the lumen and carrying matrix-dense particles whose barrel shape and faceted ends identify them as otoconia (Figure 4, D, F, and F′). Roof-associated otoconia tended to be smaller than macular otoconia, although with the few particles available from a single saccule, the difference did not reach significance (P = 0.151; Figure 4, G and G′); it is established quantitatively in the human utricle below (Figure 5F). Multiplex labeling placed Oc90 and dentin matrix protein 1 (DMP1) in clear vesicles within the RARA epithelium and in the matrix outside it (Figure 4H). A 10-kDa anionic, membrane-impermeant fluorescent dextran (27) labeled the same compartments, including vesicle clusters containing otoconial particles. Dextran labeling was abolished by decalcification (mouse saccule, Supplemental Figure 2), indicating that it reports mineral rather than matrix; crystal and cargo protein thus occur together on both sides of the apical membrane. In the RARA, therefore, crystallizing domains lie within roof epithelial cells, otoconia-forming proteins occur together with mineral in intracellular vesicles, and small, nascent otoconia occupy the ECM strands leaving the apical surface.

**Figure 4.**
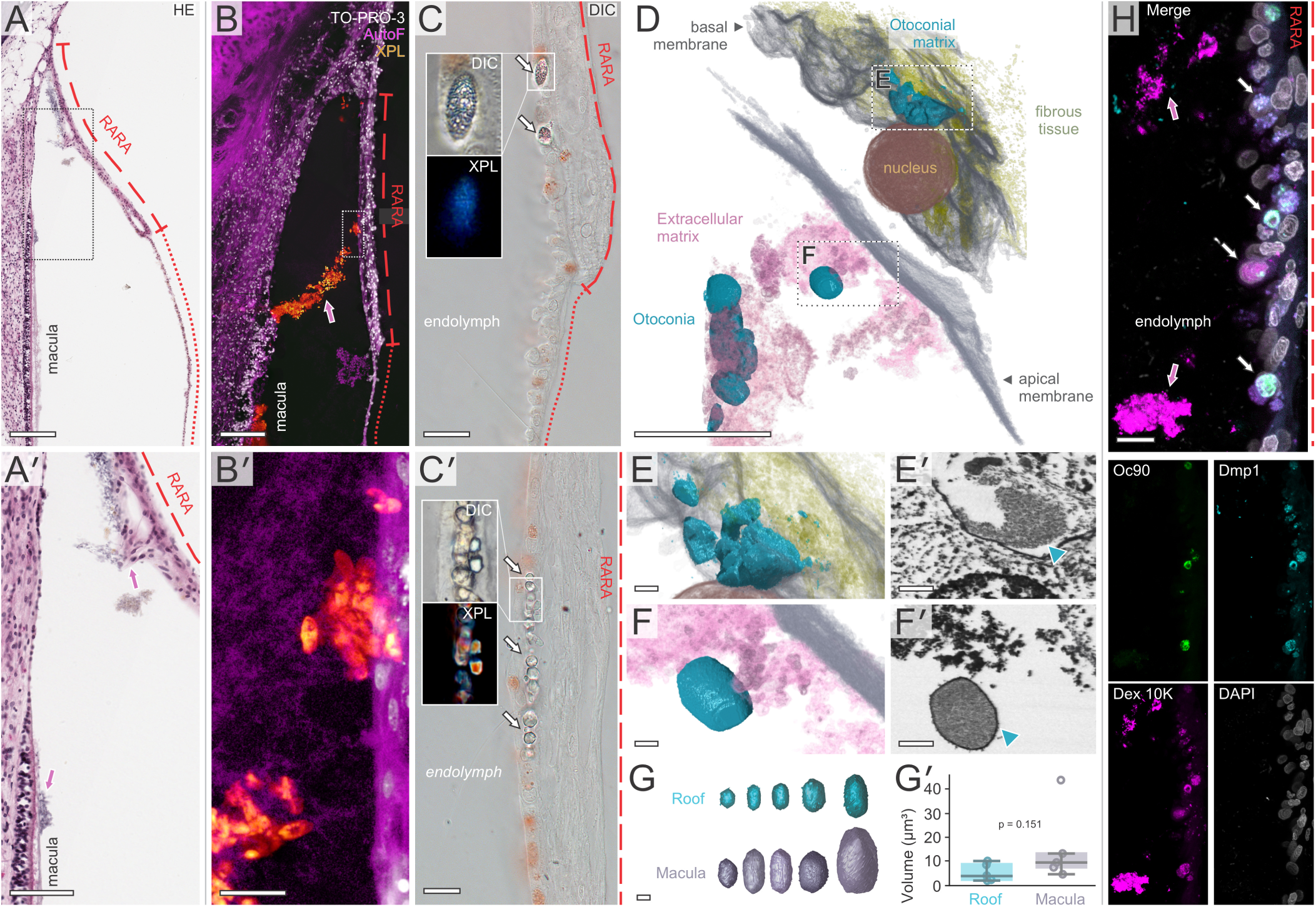
The human saccular roof domain forms an otoconial conduit and shows features of active otoconia biogenesis. (**A** and **A′**) Decalcified, celloidin-embedded saccule, hematoxylin and eosin (H&E): roof (dotted line) with its reinforced anterior roof area (RARA, dashed line); box enlarged in **A**′: otoconial protein remnants (magenta arrow) on the RARA and the anterior macular margin. (**B** and **B′**) Mineralized, Technovit-embedded saccule, cross-polarized light (XPL) with autofluorescence (AutoF; magenta) and TO-PRO-3 nuclear stain (white): birefringent otoconia (magenta arrow) along the conduit from RARA to anterior macular margin; box enlarged in **B**′: otoconia within the autofluorescent scaffold extending from the roof. (**C** and **C′**) Differential interference contrast (DIC), mineralized RARA: ovoid, weakly birefringent particles in the epithelium and subepithelial space (arrows, **C**) and birefringent otoconia in the subepithelial space (arrowheads, **C′**); insets, DIC and XPL of the marked particles. (**D**–**G′**) Focused ion beam–scanning electron microscopy (FIB-SEM), RARA. (**D**) 3D reconstruction: otoconial matrix (cyan) near the nucleus; otoconia (teal) in extracellular matrix (pink) beyond the apical membrane; boxes, regions in **E** and **F**. (**E** and **E′**) Perinuclear matrix-dense material within a roof epithelial cell (arrowhead, **E′**). (**F** and **F′**) Barrel-shaped otoconium in extracellular matrix near the apical membrane (arrowhead, **F′**). (**G** and **G′**) Five reconstructed roof-associated and five macular otoconia (**G**) and their volumes (**G′**; circles, individual otoconia; boxes, median and interquartile range; whiskers, 1.5 × interquartile range; P = 0.151, 2-sided Mann–Whitney U test). (**H**) RARA immunofluorescence, mineralized section: otoconin-90 (Oc90; green), dentin matrix protein 1 (DMP1; cyan), 10-kDa anionic dextran (Dx 10K; magenta), DAPI (white): intracellular vesicles positive for Oc90, DMP1, and Dx 10K (black arrows); strongly Dx 10K–positive scaffold material in the lumen (magenta arrows), with Oc90- and DMP1-positive particles. Scale bars: 500 μm (A); 200 μm (B); 20 μm (B′, C, and C′); 10 μm (D and H); 1 μm (E, E′, F, F′, and G).

**Figure 5.**
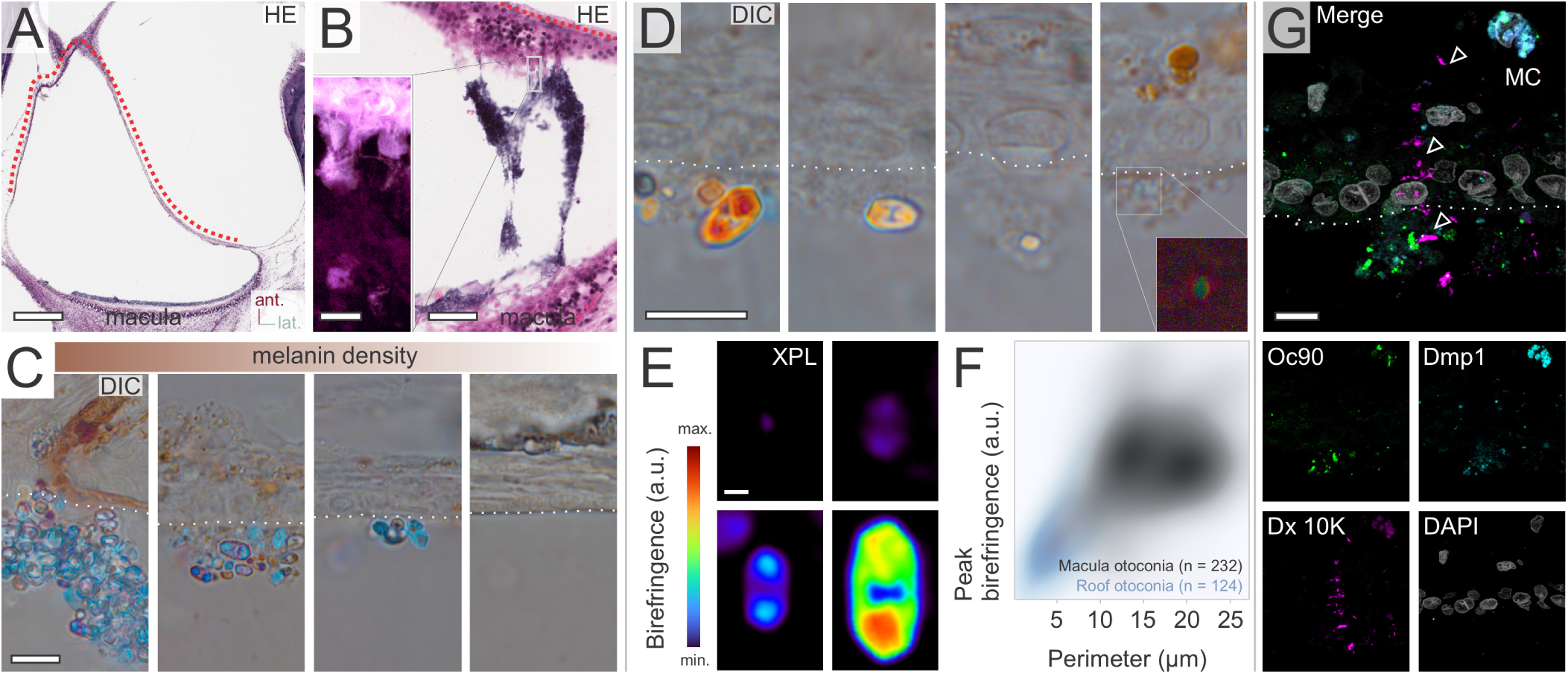
The human utricular vestibular dark cell (VDC) roof domain forms an extracellular matrix (ECM) web containing nascent otoconia. (**A**) Decalcified, celloidin-embedded utricle, hematoxylin and eosin (H&E): roof (dotted line) opposite the macula; ant., anterior; lat., lateral. (**B**) Same preparation: web-like scaffold connecting roof (dotted line) and macula; inset, boxed region by confocal eosin fluorescence, bleb-like protrusions at the epithelial surface. (**C**) Mineralized, Technovit-embedded roof, differential interference contrast (DIC), fields ordered by local melanin density (gradient bar): grape-like packets of birefringent otoconia at the VDC surface, packets and otoconia larger where melanin-containing cells are abundant and smaller as subepithelial melanin declines outside the VDC area. (**D**) DIC: otoconia of varied size at the VDC surface, down to weakly birefringent submicrometer particles within material attached to the apical epithelial surface (inset, cross-polarized light [XPL]). (**E**) XPL with birefringence heat map (color scale, arbitrary units [a.u.]) of VDC-associated otoconia: from weakly birefringent submicrometer particles to highly birefringent barrel-shaped otoconia with hexagonal ends. (**F**) Peak birefringence versus perimeter, macular (gray, n = 232) and VDC-associated (blue, n = 124) otoconia from 2 utricles: VDC-associated otoconia significantly smaller and less birefringent (2-sided Kolmogorov–Smirnov test: perimeter, D = 0.74, P < 0.0001; peak birefringence, D = 0.54, P < 0.0001). (**G**) Mineralized section, immunofluorescence: otoconin-90 (Oc90; green), dentin matrix protein 1 (DMP1; cyan), 10-kDa anionic dextran (Dx 10K; magenta), DAPI (white): Oc90 and DMP1 in vesicles of the melanin-containing cells (MC) and epithelium and in the extracellular matrix; Dx 10K–positive vesicles extending from the subepithelial zone into the matrix (arrowheads); dotted line, epithelial surface. Scale bars: 500 μm (A); 50 μm (B); 10 μm (B, inset, C, D, and G); 1 μm (E).

The human utricular roof domain (Figure 5A) showed the same features in a different geometry, giving rise, as in the mouse, to a dense otoconial web extending toward the macula (Figure 5B). Differential interference contrast imaging showed grape-like packets of birefringent otoconia over the VDC domain, of widely varying sizes and generally smaller than macular otoconia (Figure 5C). In every specimen, packet abundance tracked the local density of non-epithelial melanin-containing cells: packets were largest and most numerous where these cells were most concentrated and diminished, then disappeared, toward the margins of the VDC area as the cells became sparse (Figure 5C). At higher resolution, the packets resolved into individual otoconia down to submicrometer particles, each embedded in extracellular material at the luminal surface of the epithelium; the smallest were identifiable only by faint birefringence, consistent with the earliest stages of crystallization (Figure 5D). Across these particles, birefringence increased with perimeter (Figure 5, E and F), and VDC-associated crystals were significantly smaller and less birefringent than macular otoconia (Figure 5F). Multiplex labeling localized Oc90 and DMP1 within the web and in vesicles of the melanin-containing cells, and dextran labeled the same structures along their course from roof domain toward macula (Figure 5G). Comparable conduits and webs were already present, and more prominent, in fetal human otolith organs at 7.5 weeks of gestation (Supplemental Figure 3), so these structures are established early in development and persist in adulthood. In both human otolith organs, therefore, the roof domain described in animals is present, contains otoconia-forming proteins, holds otoconia at successive early stages of mineralization, and is linked to the macula by ECM scaffolds carrying the same material. The least mineralized particles were confined to the roof domain and its scaffolds and mature otoconia to the macular surface, an ordering that places the origin of macular otoconia upstream of the macula itself.

### Single-cell transcriptomics implicates a specialized roof mesenchyme, signaling to the roof epithelium, in otoconia and scaffold production

In every roof domain examined here, the specialized roof epithelium overlies a distinct subepithelial cell population, which we term the roof mesenchyme. This mesenchymal layer is prominently seen in a pigmented region beneath the VDC epithelium of the utricle and as an unpigmented region beneath the RARA of the saccule (Figures 4 and 5; Supplemental Figure 4). In the human utricle, otoconial packet abundance tracked the density of this melanin-containing mesenchyme (Figure 5C), and the same layer lies directly beneath the roof epithelium of the postnatal day 4 (P4) mouse utricles (Figure 6A), suggesting that the roof domain operates as an epithelial-mesenchymal niche rather than just as an epithelial surface. To test which transcriptional programs and signals underlie this specialized cellular region, we reanalyzed a previously published neonatal mouse utricle scRNA-seq dataset (28). Cell clusters enriched for otolith mineralization-related gene modules (GO:0045299: *Atg4b*, *Atp2b2*, *Oc90*, *Otol1*, and *Otop1*) were found in the roof epithelial cells, transitional epithelial cells, macular supporting cells, and mesenchymal cells (Figure 6, B and C). After unsupervised re-clustering, three epithelial and two mesenchymal subpopulations were resolved (Figure 6D). By inferring cell-cell communication from ligand and receptor gene expression, mesenchyme-to-epithelium signaling was ranked as the dominant axis among these populations, with ECM-receptor interactions being especially prominent (Figure 6E). The most notable interactions were between mesenchymal collagen ligands engaging with syndecan 4 (Sdc4) and integrin receptors on the epithelial populations (Figure 6F; the individual ligand-receptor pairs of the collagen pathway, the top-ranked signaling pathway, are shown in Supplemental Figure 5). To relate the identified subclusters to the tissue, we used annexin A2 (Anxa2) and cochlin (Coch), whose expression differed between the epithelial and mesenchymal clusters (Figure 6G). Morphologically specialized mesenchymal cells lined the perilymphatic side of the utricular roof epithelium but were absent along the adjoining ampullary roof (Figure 6H), and immunofluorescence showed a thick Anxa2-positive mesenchymal layer beneath the utricular roof epithelium, whereas the ampullary roof carried only a thin, Coch-positive mesenchyme without Anxa2 (Figure 6, I–I′′). This Anxa2-positive layer likely corresponds to the melanin-containing roof mesenchyme whose abundance predicts the otoconial material above it.

**Figure 6.**
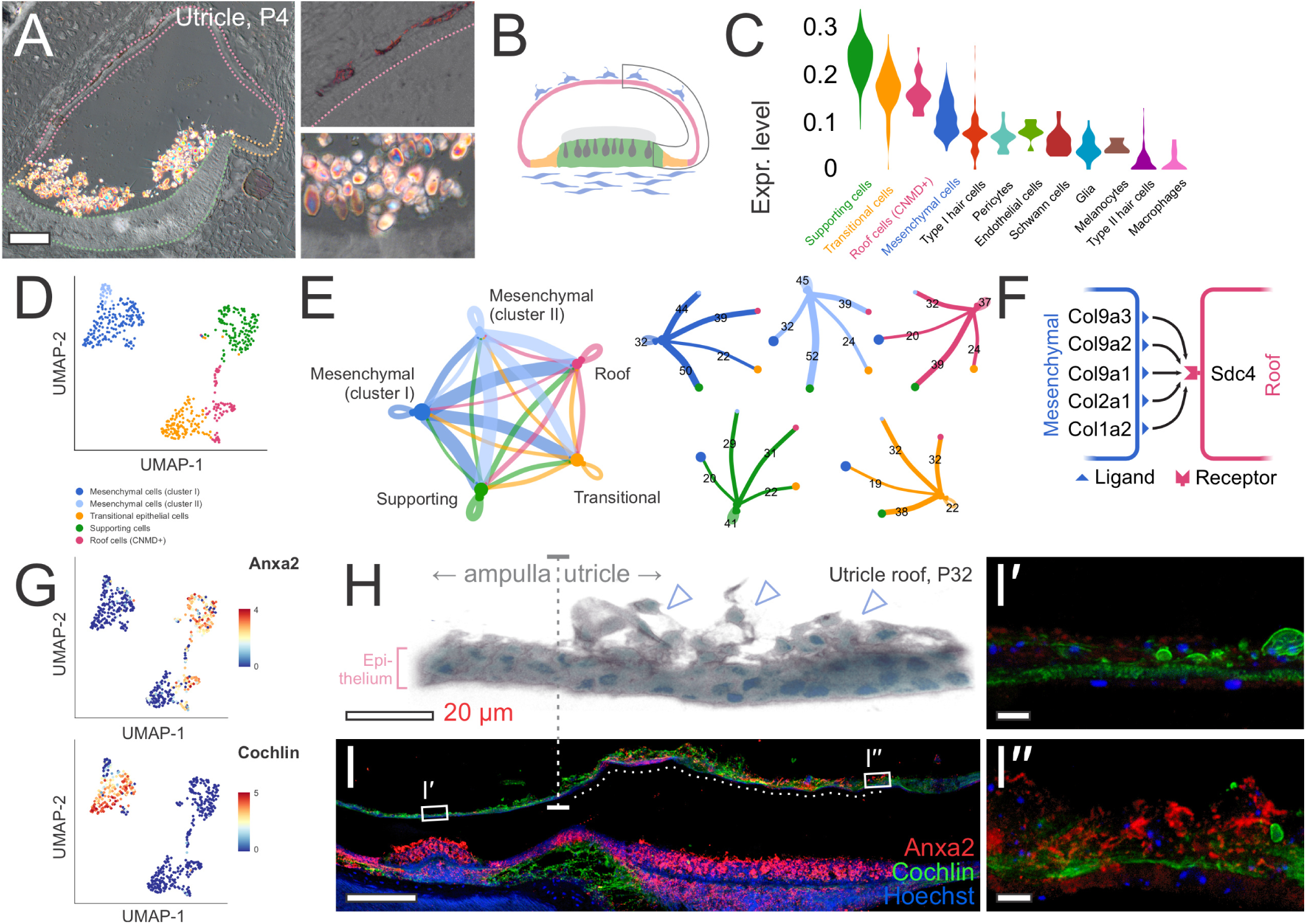
Roof epithelial-mesenchymal signaling programs implicated in scaffold assembly and otoconia biogenesis. (**A**) Mineralized, Technovit-embedded postnatal day 4 (P4) mouse utricle, polarized light: birefringent otoconia over the macula; roof epithelium outlined in magenta, transitional epithelium in orange, macular epithelium in green; insets, melanin-containing cells beneath the roof epithelium (top) and birefringent otoconia (bottom). (**B**) Schematic of the utricular cell types in **A**: roof epithelium (magenta), transitional epithelium (orange), macular epithelium (green), and mesenchymal cells (blue). (**C**) Violin plots of otolith mineralization (GO:0045299) gene module expression across all cell types of the P4–P6 mouse utricle single-cell RNA-sequencing (scRNA-seq) dataset: highest in the supporting, transitional, roof, and mesenchymal populations analyzed further (colored labels). (**D**) Uniform manifold approximation and projection (UMAP) of the subsetted data after re-clustering: roof cells (chondromodulin positive, *Cnmd*+), transitional epithelial cells, supporting cells, and mesenchymal cells, the last resolved into two subclusters (clusters I and II). (**E**) Cell-cell communication analysis: aggregated network (left) and per-population networks (right), with the two mesenchymal clusters the dominant senders to all other populations. (**F**) Predicted collagen–syndecan 4 (Sdc4) signaling route from the mesenchymal clusters (ligands Col1a2, Col2a1, Col9a1, Col9a2, and Col9a3) to the roof epithelium. (**G**) UMAP feature plots of Anxa2 and Coch (cochlin), the markers used in **I**–**I′′**. (**H**) Confocal autofluorescence, P32 utricular roof at the ampulla–utricle junction (vertical dashed line), epithelium bracketed: morphologically specialized mesenchymal cells (arrowheads) on the perilymphatic side of the utricular roof, absent along the ampullary roof. (**I**–**I′′**) Annexin A2 (Anxa2; red), cochlin (green), and Hoechst 33258 (blue) immunofluorescence: low-magnification overview of the utricle and the adjoining horizontal semicircular canal ampulla (**I**; vertical dashed line, ampulla–utricle junction; dotted line, utricular roof epithelium; boxes, regions in **I′** and **I′′**): ampullary roof with a thin, cochlin-positive mesenchyme (**I′**); utricular roof with a thick, strongly Anxa2-positive mesenchymal layer (**I′′**), showing that the Anxa2-positive specialized mesenchyme is confined to the utricular roof. Scale bars: 100 μm (**A**); 20 μm (**H**, **I′**, and **I′′**); 200 μm (**I**).

### The two-layered roof domain and its scaffold are conserved across vertebrates

The roof-macula arrangement was not confined to mouse and human. In archival saccules from lizard, opossum, rat, gerbil, cat, sea lion, and pig, a continuous ECM conduit extended from the non-sensory roof domain toward the macular margin, cable-like in lizard and more reticular in mammals, with species-dependent variation in thickness and branching (Supplemental Figure 4). A morphologically distinct roof mesenchyme was identifiable at its origin in most species, consistent with conservation of the two-layered roof domain together with its scaffold. The utricular web was too inconsistently preserved in this archival material for comparable analysis. Scaffold-based coupling of roof domain and macula is therefore a general feature of otolith organ organization rather than a mammalian specialization, present in a squamate and across marsupial and placental mammals.

## Discussion

In BPPV, the most common form of vertigo, dislodged otoconia enter a semicircular canal and trigger positional vertigo (5–7, 13). With age, otoconial mass declines (16, 17), blunting gravity sensation (12), predisposing to imbalance and falls (1–4). Here we identify, in human and animal otolith organs, the cellular niches in which otoconia are formed, characterize them at the structural, ultrastructural, and transcriptional level, and reveal previously unrecognized matrix scaffolds that likely guide otoconia from these niches to the macula; from this anatomy we derive testable cell- and structure-based models of the failure points that may underlie both disorders.

### The roof epithelial-mesenchymal niche, not the macula, is the primary site of otoconia formation

Studies in mice, zebrafish, and birds had shown that otoconial proteins are produced not only by the macula but also by the non-sensory roof epithelium of the otolith organs (18–21), yet their assembly into crystals was still assumed to occur on the macular surface. This assumption is disputed by the present data, which place the onset of mineralization in the roof domain. In mouse and human otolith organs alike, mineralized otoconia were present at the roof. In the human saccule, otoconia ranged from intraepithelial crystalline particles to fully formed crystals within the niche but were overall smaller and less mineralized than those of the macular layer, suggesting that nascent otoconia originate in the roof domain. The roof niche itself consists of two interacting cell layers: the roof epithelium and a distinct, region-restricted mesenchyme of larger cells beneath it. In the utricle, this mesenchyme is pigmented and lies under the VDC epithelium; in the saccule, it is unpigmented and lies under the RARA. Transcriptomic subclustering of the neonatal mouse utricle identified this mesenchyme as the dominant signaling source to the roof epithelium, principally through collagen-Sdc4 and collagen-integrin signaling, nominating it as a candidate organizer of the ECM programs that equip the roof epithelium to produce otoconial matrix and scaffold. In tissue, the utricular roof mesenchyme was strongly immunoreactive for the secretion-associated protein Anxa2, which was absent from the ampullary roof (Figure 6). Although the dataset is from an early postnatal time point rather than from adult mice, the signaling programs it nominates are active well after otoconia formation begins in the mouse (21), and nascent otoconia persist in the roof domains of adult mice and humans. This is consistent with a program that remains active in adult life.

### Matrix scaffolds connect the roof niche to the macula and are implicated in maintaining the macular otoconial mass in adult life

Formation at the roof raises the question of how nascent otoconia, already mineralized and therefore subject to gravity, reach the macular surface, where their mass loads the sensory receptors. In both otolith organs we found, unexpectedly, a roof-derived matrix structure—a slender conduit in the saccule, a diffuse web in the utricle—that spans the fluid-filled lumen, is glycoprotein and fibronectin rich, and carries Oc90-positive nascent otoconial packets into the macular otoconial layer, in the human organs as in the mouse. That such consistent structures went unrecognized could be explained by prior technical limitations: they are filigree, collapsed or dissolved by routine decalcification and histological processing, and their micrometer-scale nascent otoconia are not resolved without crystal-sensitive imaging. Two features indicate that they are functional routes rather than developmental remnants. First, macular otoconia grade in size away from the point of scaffold contact, consistent with continuous seeding there and maturation on the macula. Second, when the scaffolds ruptured in adult guinea pigs, the otoconial layer decreased in thickness and was eventually lost. Notably, this structural arrangement has a precedent in the cochlea (the inner ear’s hearing organ), where the tectorial membrane, which couples sound-induced motion to the hair cell bundles, is assembled from tectorins and type II collagen secreted apically by the non-sensory interdental cells and remains anchored to them and is replenished throughout life (29). In the otolith organs, the roof niches may export matrix in the same way, except that theirs carries mineralized otoconia. While real-time movement along the scaffolds was not observed, its direction and progression were inferred from the gradient of mineralization.

### The otoconia life cycle predicts novel failure points in common vestibular disorders

Our data suggest a life cycle of otoconia with structural stations across the entire otolith organ where otoconial seeds begin at the roof niche, are transported along the scaffold, and mature within the macular layer (Figure 7, A and B). If this cycle runs continuously in adult life, as the data indicate but future studies must confirm, it implies a fourth station, clearance: new otoconia arriving continuously must be matched by removal of old ones at a similar rate for the otoconial mass and macular receptor loading to stay constant. We found no structural sign of such clearance, and where it occurs remains open. Each station, however, is a potential failure point, and none is represented in current disease models (13–17, 30, 31):

**Figure 7.**
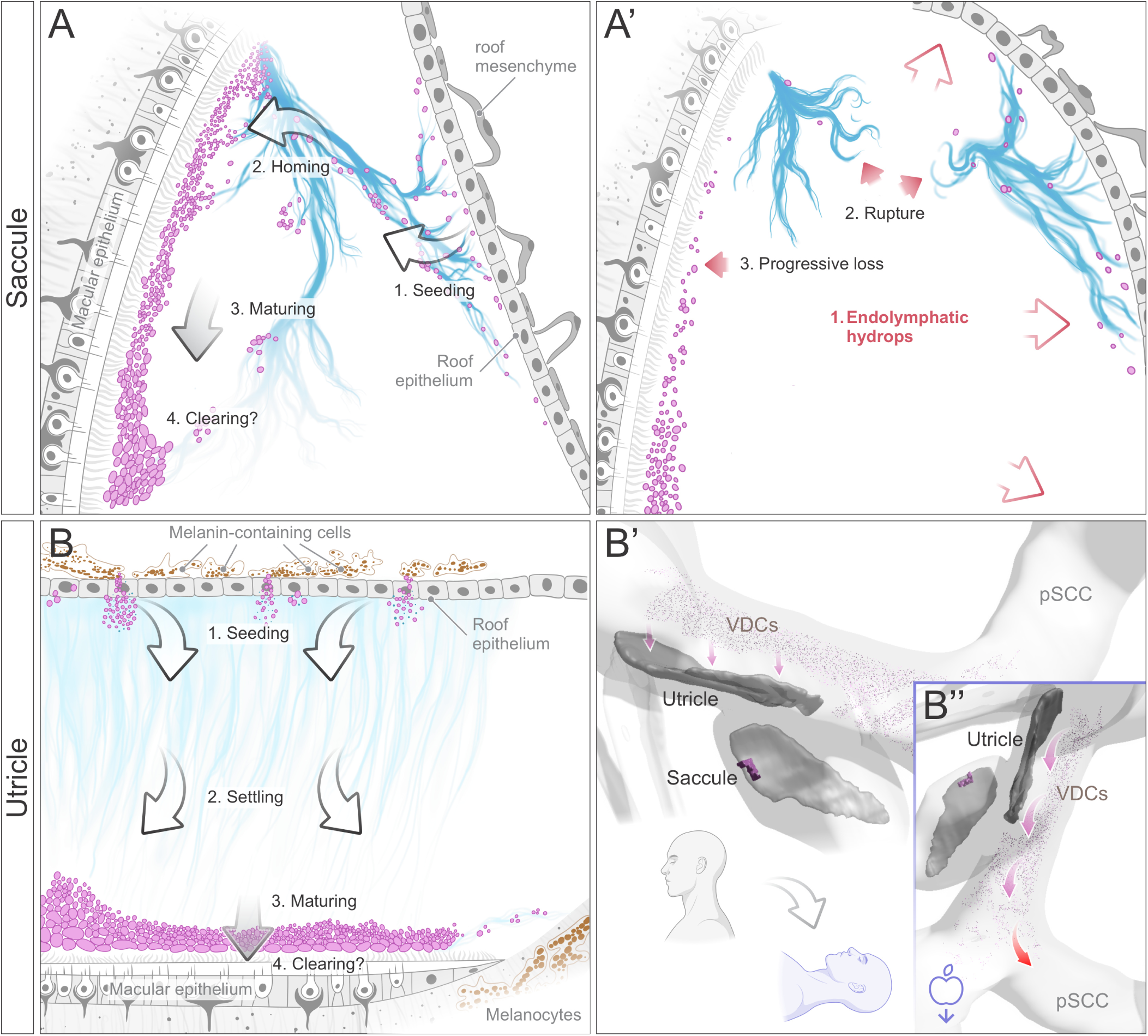
Conceptual model of roof-derived otoconia biogenesis, scaffold-guided macular delivery, and their failure in endolymphatic hydrops and benign paroxysmal positional vertigo (BPPV). Schematics, not to scale. (**A**) Saccule: otoconial protein and nascent otoconia (magenta) from the roof epithelial-mesenchymal niche are seeded onto the extracellular matrix (ECM) conduit (blue) (1, seeding), travel to the anterior macular insertion (2, homing), migrate across the macula while enlarging (3, maturing), and are removed in a hypothesized final step for which we saw no morphological evidence (4, clearing; question mark). (**A′**) Endolymphatic hydrops (1): distension (open arrows) moves the roof away from the macula and ruptures the conduit (2), with progressive loss of the macular otoconial layer (3), as observed in the guinea pig (Figure 3). (**B**) Utricle: the roof epithelium and underlying melanin-containing cells (MC) of the roof mesenchyme seed otoconial protein and nascent otoconia into the diffuse ECM web (1); particles settle along the gravity vector onto the macula (2), mature in a depth gradient within the otoconial layer (3), and are hypothetically cleared (4; question mark). (**B′** and **B′′**) Proposed BPPV mechanism in a 3D model of the human vestibule with utricular and saccular maculae. Upright (**B′**): otoconia from the vestibular dark cell (VDC) field settle along the gravity vector onto the utricular macula (arrows; figures, head position). Recumbent (**B′′**): the gravity vector drives otoconia still in the web toward the posterior semicircular canal (pSCC; red arrow; icon, gravity direction), so material released from the web would cause canalithiasis.

In BPPV, the standard model explains the symptoms but not their origin: mature otoconia detach from the utricular macula (13–15) and, if loose in a semicircular canal, can turn head movements into false signals of rotation and brief spells of vertigo. However, why or where the dislodged otoconia originate is not part of the model. Our data point to the utricular web as a novel structural failure point. The web is the route by which nascent otoconia reach the utricular macula, and it appeared more diffuse and delicate than the saccular conduit. Whether its cargo arrives may depend on head position. In the upright head, the near-horizontal human macula (32, 33) lies below its roof domain, and nascent otoconia in the web settle onto it along the gravity vector. When the head is recumbent, for hours every night, otoconia midway must be held by the web, or gravity draws them toward the posterior semicircular canal, the canal most often affected in BPPV (7, 8). A fragile web may therefore shed its cargo during sleep, which would fit the clinical pattern: attacks are typically provoked by turning in bed or sitting up in the morning (8), and the affected ear tracks the habitual sleep position (34) (Figure 7, B′ and B′′). This raises two testable hypotheses about BPPV. The particles that trigger it may be nascent otoconia shed from the roof domain or the utricular web—never fully anchored and thus prone to detachment—rather than mature otoconia dislodged from the macula. The web itself may grow more fragile with age, as the niche secretes less matrix or its existing protein degrades. Both of these hypotheses could be tested by examining the roof domain and web in postmortem human otolith organs spanning the age range, as well as in tissue from patients with recurrent BPPV.

During age-related loss of macular otoconia (16, 17, 30), where there is less otoconial mass and decreased receptor loading, the organ is less capable of responding to gravity and linear acceleration (12). This weaker input for the control of posture and movement therefore makes imbalance and falls more likely (1). The current conceptual model attributes this loss to degeneration occurring in place. However, if the adult otoconial layer depends on a continuous supply as our data might suggest, the age-related decline of niche output could limit the supply in both otolith organs while the macula continues to lose crystals, and part of the loss would be a failure of supply. To test this, the adult niche can be profiled in animals for whether its program persists and declines with age, and its output and scaffold integrity measured in the same temporal bone series.

The third failure point is the one we could observe directly. Endolymphatic hydrops is characterized by the expansion of the fluid-filled lumen, common across chronic inner ear disorders and defining in Meniere’s disease (23). During this expansion, the roof and macular epithelia grow farther apart, straining the coupling provided by the otoconial trafficking scaffolds between them. In our guinea pig data, as the lumen expanded the scaffold stretched and tore and the otoconial layer began to decline as it ruptured. Two earlier observations in this model fit that sequence. First, endolymph calcium rises over the same interval (35), which the disintegration of macular otoconia we saw at 8 weeks could feed. Second, degraded otoliths accumulate on the VDC area over altered pigmented cells (36), which we would reinterpret as nascent otoconia stranded at the roof domain once the web that guides them to the macula is gone. Uncoupling of the roof and macular epithelia (Figure 7A′) may thus be a direct structural sequela of hydrops, one that depletes the macula and displaces otoconia into the lumen. This is consistent with the increased prevalence of BPPV in those diagnosed with Meniere’s disease (37–39) and in other hydropic disorders (40, 41). Because duct obstruction is a broad insult that changes endolymph volume and composition together, the guinea pig data establish a temporal association, not a cause; the decisive test is to disrupt the scaffolds selectively, without hydrops, by targeting defined scaffold components or the mesenchymal signals that induce them in transgenic models. If that reproduces the loss, the weeks between distension and rupture would define a window in which limiting hydrops protects the otoconial mass, a clinical target upstream of the symptoms.

## Methods

### Sex as a biological variable

Mouse studies included male and female animals, and comparable findings were obtained in both sexes. Guinea pig and human temporal bone cohorts included specimens from both sexes. The roof domains and scaffolds examined here are not known to be sexually dimorphic, and the findings are expected to apply to both sexes.

### Animals and tissue collection

Adult CBA/CaJ mice (Jackson Laboratory, stock 000654; 6–10 weeks of age; both sexes) were used for histologic and imaging analyses. Animals were euthanized with ketamine/xylazine followed by decapitation, and vestibular organs were harvested immediately and processed using tissue-preserving workflows that maintain the endolymph-facing architecture of the utricle and saccule (maculae, roof domains, luminal ECM, and otoconia). Specimens were embedded either in celloidin after decalcification, for histology and serial-section reconstruction, or mineralized in methyl methacrylate resin (Technovit), for imaging of otoconia in situ. CBA/CaJ mice at postnatal day 4 (P4) and P32 were used for the roof domain analyses in Figure 6. For pathology-associated analyses, we evaluated archival guinea pig inner ears subjected to uni- or bilateral endolymphatic duct obstruction at 6–8 weeks of age with survival times of 2, 8, or 16 weeks; these had been processed with the same decalcified celloidin workflow and permanently coverslipped after staining. Unoperated ears from the same cohort served as controls.

### Human temporal bones and microdissected saccule

Postmortem human temporal bones (n = 10; age range 22–83 years; 3 male, 2 female, sex not recorded for the remaining 5), together with an additional fetal specimen (7.5 weeks of gestation, male), were from the temporal bone collections of Mass Eye and Ear and the University Hospital Zurich (Zurich, Switzerland). For FIB-SEM, one fresh human saccule was microdissected from a donor specimen (woman, aged 60–65 years; postmortem interval 53 hours) under isotonic conditions, preserving the endolymph-facing epithelial architecture as far as possible.

### Histochemistry

Hematoxylin and eosin (H&E) staining was used for overall anatomy and the relationship of roof domains to the maculae. PAS-Alcian blue staining labeled glycoprotein- and proteoglycan-rich ECM structures spanning the lumen, including roof-derived conduits and webs and otoconial matrix remnants. H&E staining of celloidin sections was performed free-floating (Harris hematoxylin, 5 minutes; 0.3% acid alcohol; 0.2% ammonia water; eosin Y, 1 minute). PAS-Alcian blue staining was performed on de-celloidinized or deacrylated sections (Alcian blue, pH 2.5, 30– 90 minutes; 1% periodic acid, 10 minutes; Schiff reagent, 10 minutes). Sections were dehydrated, cleared, and mounted in Permount.

### Immunofluorescence labeling

Mouse decalcified celloidin sections were labeled for otoconin-90 and fibronectin (otoconial cargo and ECM scaffolds) and, where indicated, for annexin A2 and cochlin (roof-adjacent cellular and matrix compartments). Human Technovit sections were labeled for otoconin-90 and dentin matrix protein 1 and, where indicated, coincubated with 10-kDa fluorescent dextran to visualize extracellular pathways in roof-adjacent luminal compartments. Mouse sections were de-celloidinized in saturated sodium methoxide (1:2 in methanol), rehydrated, and blocked with 5% normal horse serum. Otoconin-90 and fibronectin were detected with rabbit anti–otoconin-90 (1:100; gift of Y.W. Lundberg) and rabbit anti-fibronectin (1:100; Abcam, ab2413), incubated overnight at room temperature, followed by goat anti-rabbit Alexa Fluor 488 (Jackson ImmunoResearch, 111-545-003) for 1 hour. For annexin A2 and cochlin, heat-induced antigen retrieval in sodium citrate buffer (pH 6.0) preceded incubation for 48 hours at 4°C with rabbit anti-ANXA2 (1:100; Sigma-Aldrich, HPA046964) and rat anti-cochlin (1:100; MilliporeSigma, MABF267), followed by biotinylated goat anti-rabbit IgG (1:400; Jackson ImmunoResearch) with streptavidin–Alexa Fluor 647 (1:400; Thermo Fisher Scientific, S21374) and donkey anti-rat Alexa Fluor 488 (1:400; Thermo Fisher Scientific, A-21208). Human Technovit sections were deacrylated in 2-methoxyethyl acetate, rehydrated, blocked with 5% normal donkey serum, and incubated overnight at 4°C with sheep anti–dentin matrix protein 1 (DMP1; 1:100; R&D Systems, AF4386) and rabbit anti–otoconin-90 (1:100), followed by biotinylated donkey anti-sheep IgG (1:500; Jackson ImmunoResearch, 713-065-147) and goat anti-rabbit Alexa Fluor 488 (1:400; Jackson ImmunoResearch, 111-545-003), and then by streptavidin–Alexa Fluor 568 (1:500; Thermo Fisher Scientific, S11223) with 10-kDa anionic dextran–Alexa Fluor 647 (1:500; Thermo Fisher Scientific, D22914) for 30 minutes. Mineralized and decalcified mouse saccule sections were labeled with the same dextran for Supplemental Figure 2. Autofluorescence in melanin-rich regions was quenched with TrueVIEW (Vector Laboratories, SP-8400). Nuclei were counterstained with Hoechst 33258 (1:1500; Thermo Fisher Scientific, H3569), DAPI (Sigma-Aldrich, D9542), or TO-PRO-3, and plasma membranes with CellMask Deep Red (1:1500; Thermo Fisher Scientific, C10046), as indicated in the figure legends.

### Light microscopy and confocal imaging

Brightfield imaging was performed for histochemical stains. Polarized-light microscopy under crossed polarizers was used to visualize birefringent calcite otoconia in mineralized specimens and to measure size–birefringence relationships in human utricles. Confocal microscopy was used for immunolabeled sections and, where applicable, for intrinsic PAS and eosin fluorescence and tissue autofluorescence. Z stacks were collected at submicrometer intervals and rendered as maximum-intensity projections for morphologic mapping. For selected human specimens, lattice structured illumination microscopy provided higher-resolution imaging of dextran-labeled extracellular pathways and roof-associated nascent otoconial material. Brightfield and polarized-light images were acquired on a Nikon Eclipse E800 or Leica DM5500 B microscope with Nikon DS-Ri2 or Leica DMC5400 cameras, and extended-depth-of-field images were merged in Zerene Stacker (v1.04). Confocal images were acquired on a Leica TCS SP8 (HC PL APO 63×/1.4 NA oil objective; 0.3-μm Z steps) and projected in Leica LAS X (v4.8). Lattice SIM was performed on a Zeiss Elyra 7 (Plan-Apochromat 63×/1.4 NA oil objective; 0.12-μm Z steps) with SIM2 reconstruction in ZEN Black.

### Focused ion beam–scanning electron microscopy

A fresh human saccule was heavy-metal stained and embedded in Araldite. Regions containing both roof domain and macular epithelium were localized on semithin toluidine blue sections, trimmed, and mounted for FIB-SEM. Two regions of interest, one in the RARA and one in the macula, were imaged on a ZEISS Crossbeam FIB-SEM (Carl Zeiss Microscopy) at the Harvard University Center for Nanoscale Systems by automated serial milling (20-nm slice thickness) and backscattered-electron imaging at pixel sizes of 7.5 nm (RARA) and 5 nm (macula). Image stacks were aligned and reconstructed in Atlas 5 (Carl Zeiss Microscopy) and Dragonfly to visualize and segment roof epithelial architecture, luminal ECM, and otoconial packets.

### Quantification of macular otoconia gradients (mouse)

For mouse maculae, confocal stacks from PAS-stained serial sections spanning the macula were processed in Fiji (42) to enhance otoconial signal and reduce non-otoconial fluorescence. Otoconia were segmented in Dragonfly using global thresholding, and size proxies were derived from shape descriptors (area/perimeter and/or volume/surface, as indicated). Otoconial coordinates were normalized from 0 to 1 along macular axes, binned into equally spaced intervals, and displayed as en face heatmaps of mean size proxy per bin.

### Quantification of hydrops severity (guinea pig)

Hydrops severity was quantified on decalcified celloidin sections as the volume of the endolymph-filled lumen (endolymphatic space) of the saccule and utricle. The endolymph-filled lumen was outlined in Dragonfly (Object Research Systems) on serial sections at 10-section intervals, excluding the sensory epithelium and the macular otoconial layer, and endolymphatic volume was estimated from the summed areas. Volumes were expressed as a percentage of the control mean. Measurements were made by an observer blinded to survival time.

### Retardance-based quantification of human otoconia

In mineralized human utricles, polarized-light microscopy was performed with standardized illumination and exposure. Individual otoconia were manually segmented to extract perimeter and peak transmitted grayscale intensity (peak birefringence) as a surrogate for retardance (degree of biomineralization). Macular and roof domain otoconia were measured in one utricle each.

### 3D reconstruction and segmentation

For whole-organ context, every tenth H&E-stained mouse inner ear section was digitized, registered, and segmented in Dragonfly to reconstruct endolymph-facing epithelia and roof domain landmarks. For composite mapping, the conduit was traced in H&E sections of 26 saccules, registered to a common saccular outline, and displayed as a kernel density estimate of tracing frequency. For FIB-SEM volumes, otoconial matrix was segmented with a supervised U-Net model as described below (Machine-learning segmentation).

### Machine-learning segmentation

Both FIB-SEM image stacks were imported into Dragonfly (Object Research Systems; v2024.1). A U-Net convolutional neural network was trained on manually annotated ground-truth labels of otoconial matrix within 16 representative slices of otoconia in the macular stack. Training used the Adam optimizer (learning rate 0.001) with a data augmentation factor of 5, including flips, rotation, zoom, and shear; an 80:20 training/validation split; and a batch size of 32, and ran for 92 epochs, with early stopping when the validation loss plateaued for 15 epochs. The resulting model achieved an average Dice coefficient of 0.75 on the validation set. The trained network was then applied to the full macular and RARA volumes, and the resulting segmentation masks were refined by 3D morphological smoothing (kernel size 10) to remove isolated noise and by manual correction as needed. The final masks enabled volumetric quantification of individual otoconia and 3D surface reconstructions.

### Single-cell RNA sequencing reanalysis and niche signaling inference

We reanalyzed a published scRNA-seq dataset of mouse utricles at P4 and P6 (28), obtained from the Gene Expression Analysis Resource (gEAR; “Interaction of mesenchymal and epithelial cells in the postnatal mouse utricle”, https://umgear.org/p?l=025ab571) (43), and analyzed it using Seurat v5 (44) in R v4.4.1. Cell cluster module scores were calculated using the otolith mineralization GO term (GO:0045299) and visualized as violin plots. After subsetting for clusters enriched for otolith mineralization, cells were re-clustered in Seurat, markers were identified for epithelial and mesenchymal subclusters, and embeddings were visualized by uniform manifold approximation and projection (UMAP). To infer cell-cell communication from ligand and receptor gene expression, we used CellChat v2.1.2 (45, 46) to estimate ligand-receptor communication probabilities; cell communication visualizations were created with the CellChat package, and data processing used default parameters and statistical cut-offs.

### Statistics

Directional gradients in otoconial size were tested by simple linear regression of bin means along each normalized macular axis (SciPy linregress), with slopes and 2-sided P values reported. Distributions of otoconial perimeter and peak birefringence were compared between roof-associated and macular populations by 2-sided Kolmogorov–Smirnov test. Particle volumes from FIB-SEM reconstructions were compared between roof-associated and macular otoconia by 2-sided Mann–Whitney U test. Endolymphatic volumes across hydrops time points are reported descriptively, because group sizes (1–2 ears per time point) preclude statistical comparison. P < 0.05 was considered significant. Biological replicate definitions and sample sizes are given in the figure legends.

### Study approval

All animal procedures were approved by the Institutional Animal Care and Use Committee of Mass Eye and Ear (protocol 2024N000071). Studies of human temporal bones, including fetal specimens, were approved by the Institutional Review Board of Mass Eye and Ear (protocol 2021P001593) and the University Hospital Zurich (Zurich, Switzerland; #BASEC-2021-00820). Postmortem tissue was procured under established protocols, with written informed consent obtained from donors or their next of kin prior to donation, and all specimens were deidentified.

## Data availability

Values underlying the graphs in Figures 1J, 2K, 3B, 3G, 4G′, and 5F are provided in the Supporting Data Values file. All other data generated or analyzed in this study are included in this article and its supplemental material. The scRNA-seq dataset reanalyzed here (Figure 6 and Supplemental Figure 5) is publicly available through the Gene Expression Analysis Resource (gEAR; https://umgear.org/p?l=025ab571), as described in David et al. (28). The code used for analysis is available from the corresponding author on request.

## Author contributions

DMC and AHE conceived the study. DMC performed experiments, collected and analyzed data, prepared figures, and drafted the manuscript. RO contributed to experimental work and to specimen preparation for FIB-SEM. SK performed FIB-SEM experiments, data collection, and image analysis. APD performed the scRNA-seq reanalysis and computational analyses and contributed to manuscript revision. RZ and TAJ contributed to scRNA-seq analysis and interpretation and to manuscript revision. AAI contributed to data interpretation and manuscript revision. AHE designed and supervised the study, performed experiments, analyzed and interpreted data, wrote and revised the manuscript, and obtained funding. All authors reviewed and approved the final manuscript.

## AI disclosures

Claude Opus 4.8 and Claude Opus 5 (Anthropic) were used in January–February 2026 and August–September 2026 to assist with first drafting and formatting of the manuscript text. A U-Net convolutional neural network in Dragonfly (v2024.1; Object Research Systems) was used to segment otoconial matrix in the FIB-SEM data, as described in Methods. The authors assume full responsibility for ensuring the integrity and accuracy of AI output presented in this article.

## Acknowledgments

The authors thank David Bächinger (Department of Otorhinolaryngology, Head and Neck Surgery, University Hospital Zurich, Zurich, Switzerland) for the procurement and processing of a subset of the human temporal bone specimens used in this study, and Andrés Felipe Correa for the schematics presented in Figure 7. This work was performed in part at the Harvard University Center for Nanoscale Systems (CNS); a member of the National Nanotechnology Coordinated Infrastructure Network (NNCI), which is supported by the National Science Foundation under NSF award no. ECCS-2025158.

