## Supplemental Figures 1 to 5 for "Disruption of a structural niche for otoconia maintenance may underlie common vestibular disorders"

This file includes Supplemental Figures 1–5.

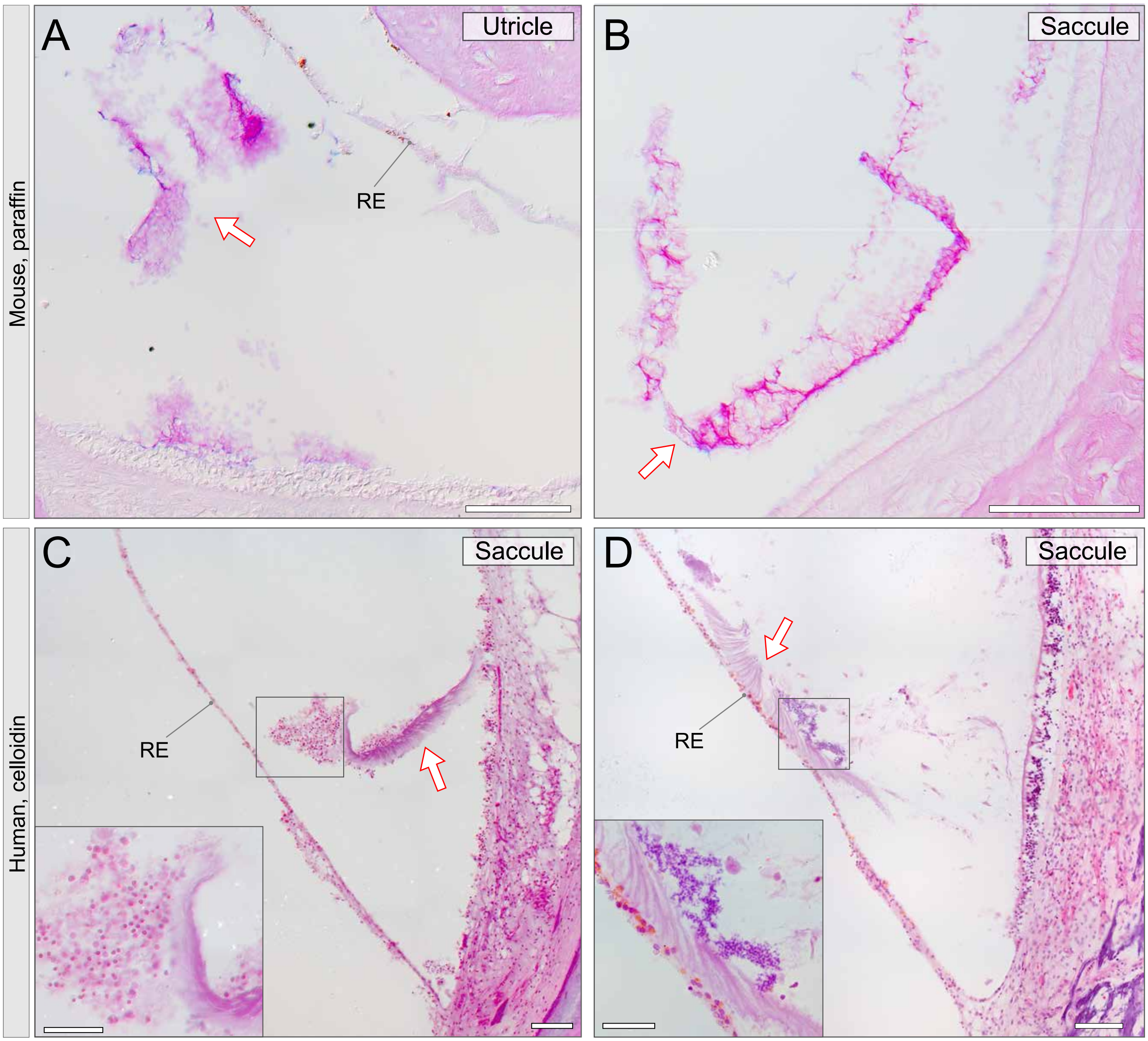


**Supplemental Figure 1. Distinguishing artifactually dislodged otoconial membranes from the saccular otoconial conduit.** Periodic acid–Schiff (PAS)–stained sections: paraffin-embedded mouse utricle (**A**) and saccule (**B**); decalcified, celloidin-embedded human saccule (**C** and **D**). Arrows, dislodged otoconial membrane floating in the lumen or attached to the roof epithelium (RE)—a processing or postmortem artifact. Unlike these loose, amorphous fragments, the otoconial conduit is a taut extracellular matrix scaffold anchored between the roof epithelium and the anterior macular rim (compare Figures 1 and 4). Scale bars: 100 μm (**A**–**D**); 50 μm (**insets**).

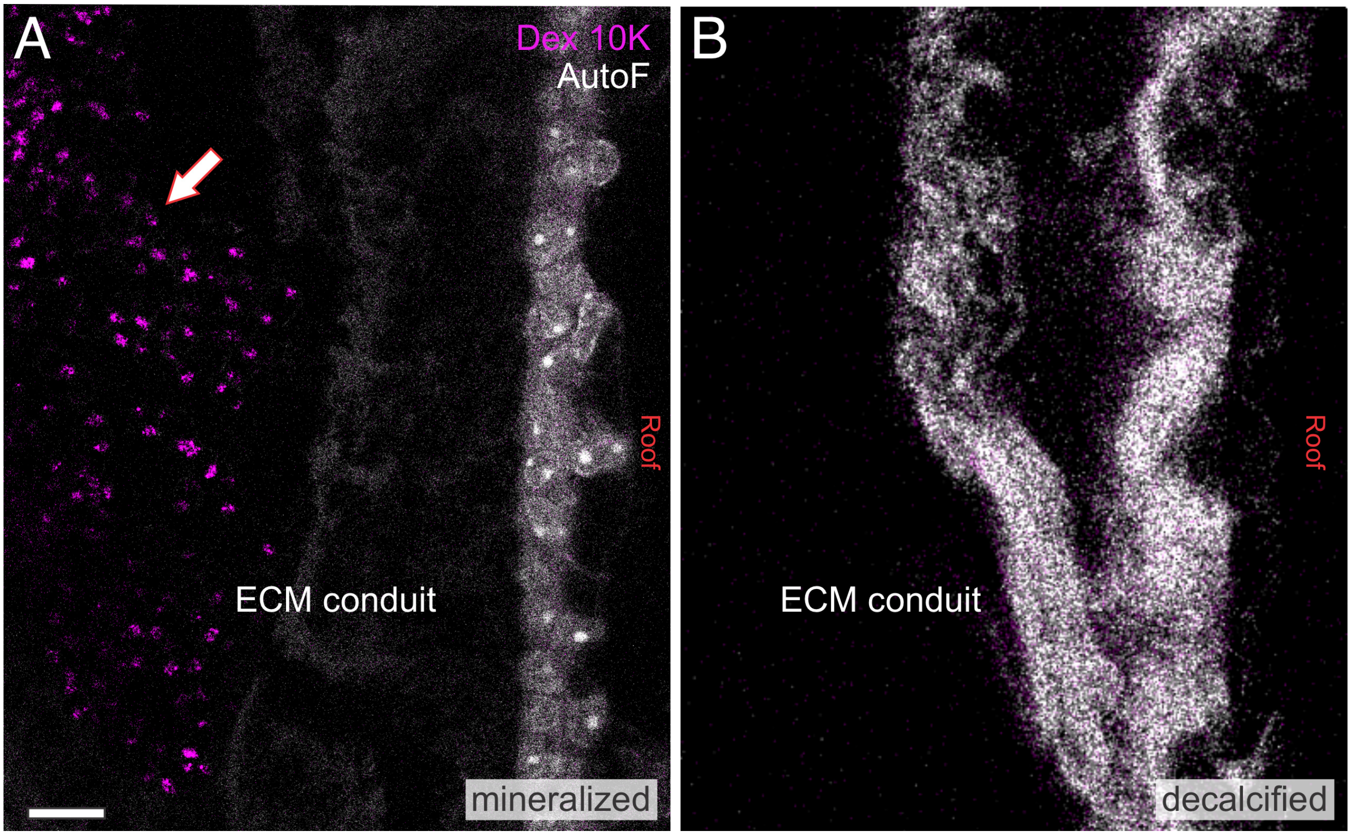


**Supplemental Figure 2. Fluorescent dextran labeling of otoconia depends on mineral.** Mouse saccule labeled with 10-kDa anionic fluorescent dextran (Dx 10K). (**A**) Mineralized, Technovit-embedded section: Dx 10K fluorescence in the otoconia (arrow) within the extracellular matrix conduit. (**B**) Decalcified section: Dx 10K fluorescence is absent from the conduit. Loss of the signal with decalcification indicates that Dx 10K labels the mineral phase rather than the organic matrix, as applied to the human specimens in Figures 4H and 5G. Scale bars: 10 µm (**A** and **B**).

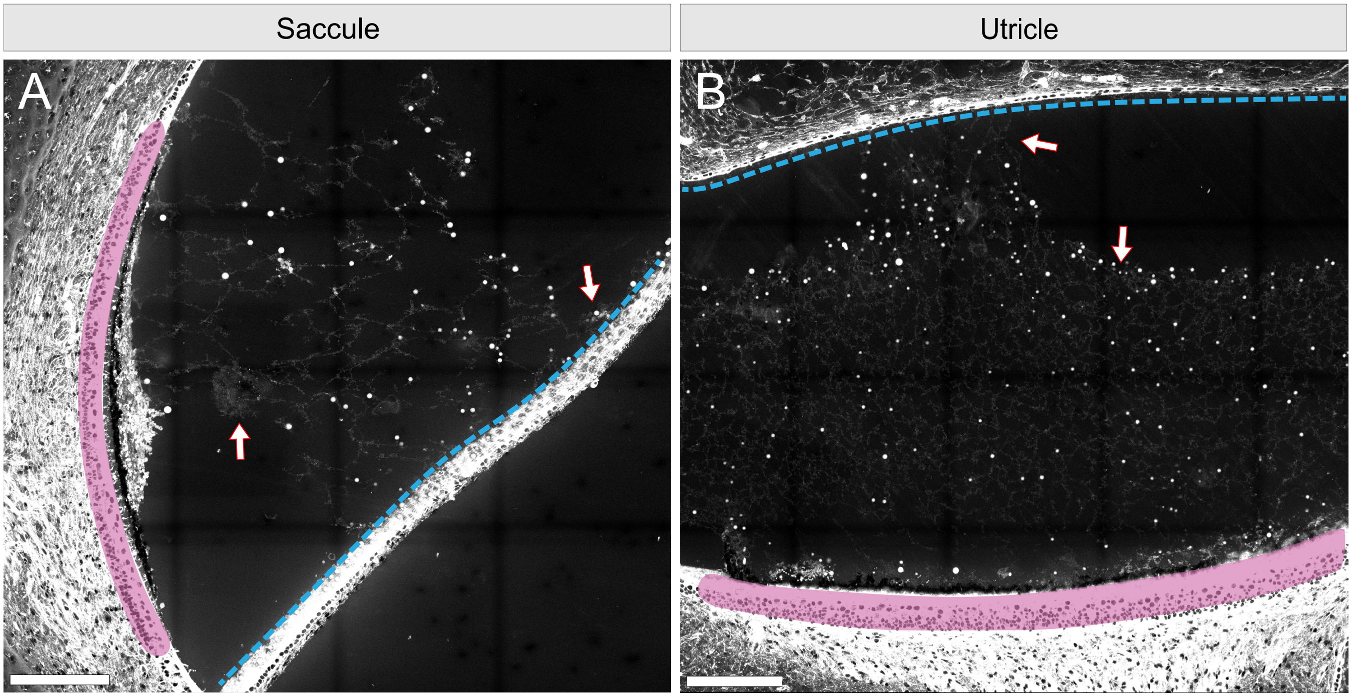


**Supplemental Figure 3. Developmental appearance of the human saccular otoconial conduit and utricular otoconial web.** Hematoxylin and eosin–stained sections at 7.5 weeks of gestation, confocal eosin fluorescence; macular sensory epithelium shaded magenta, roof epithelium traced with a blue dotted line. (**A**) Saccule: arrows, roof-derived otoconial conduit spanning the lumen toward the anterior macular rim, at this stage a matrix-rich nascent strand. (**B**) Utricle: vestibular dark cell (VDC) domain bracketed; arrows, filamentous otoconial web arising from VDC apices, abundant and widespread at this stage, with particulate material along its fibers. Scale bars: 100 μm (**A** and **B**).

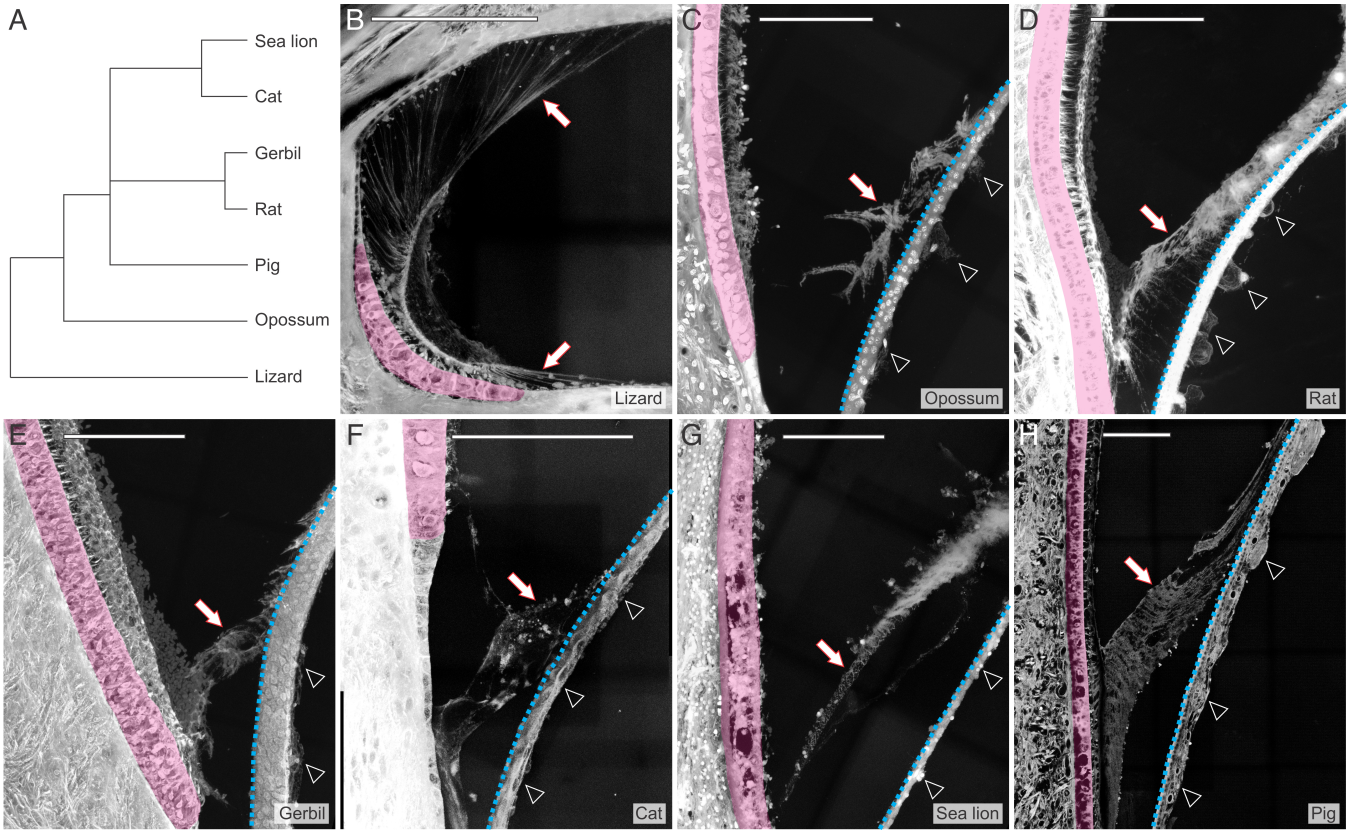


**Supplemental Figure 4. Conservation of the saccular otoconial conduit across vertebrates.** (**A**) Phylogenetic tree of the taxa examined. (**B**–**H**) Decalcified sections of lizard (**B**), opossum (**C**), rat (**D**), gerbil (**E**), cat (**F**), sea lion (**G**), and pig (**H**) saccules: macular sensory epithelium shaded magenta; roof epithelium traced with a blue dotted line; arrows, extracellular matrix conduit spanning the lumen from roof epithelium to anterior macular rim; arrowheads, roof mesenchyme at the conduit's origin. Species trend: coherent, cable-like conduit in the lizard, more reticular in mammals, continuity from roof domain to macula maintained. Shading and overlays added for clarity. Scale bars: 100 μm (**B**–**H**).

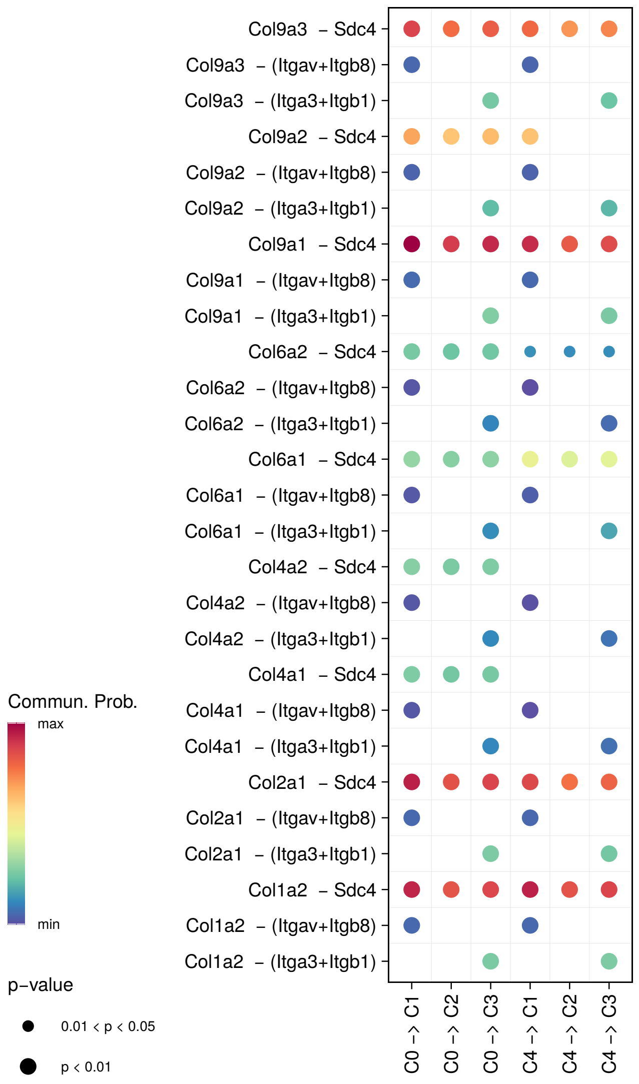


**Supplemental Figure 5. Collagen ligand-receptor interactions between utricular cell populations inferred from single-cell RNA sequencing.** CellChat ligand-receptor analysis of the subsetted and re-clustered postnatal day 4–6 (P4–P6) mouse utricle scRNA-seq data (Figure 6, D and E), showing the individual ligand-receptor pairs of the collagen pathway, the top-ranked signaling pathway by information flow, from the mesenchymal to the epithelial populations: collagens from the mesenchymal clusters engage syndecan 4 (Sdc4) and integrin heterodimers on the epithelial populations. C0, mesenchymal cells (cluster I); C1, supporting cells; C2, transitional epithelial cells; C3, roof cells (Cnmd+); C4, mesenchymal cells (cluster II). Dot color, communication probability; dot size, P value.
